## Supplementary Figures and Tables for "Genetic architecture and lifetime dynamics of inbreeding depression in a wild mammal"

Authors names and addresses:

Stoffel, M.A.[1*], Johnston, S.E.[1], Pilkington, J.G.[1], Pemberton, J.M[1]

[1]Institute of Evolutionary Biology, School of Biological Sciences, University of Edinburgh, Edinburgh, EH9 3FL, United Kingdom

* Corresponding author:
Martin A. Stoffel
Postal address: Institute of Evolutionary Biology, University of Edinburgh, Edinburgh, EH9 3FL, UK


### Supplementary Figures

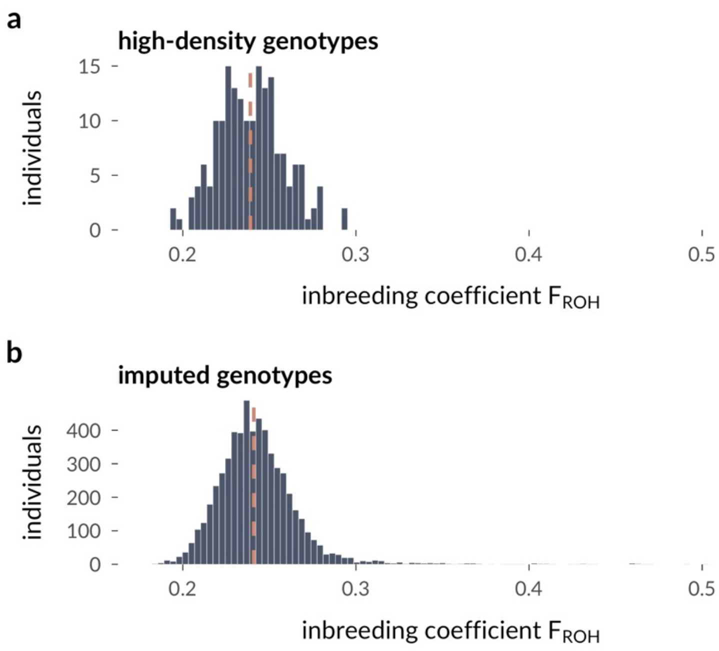


**Supplementary Figure 1: Distribution of $F_{ROH}$ for individuals with non-imputed (high-density) and imputed SNPs. a** Distribution of inbreeding coefficients $F_{ROH}$ of 181 individuals with annual survival data available who were genotyped on a high-density SNP chip. **b** $F_{ROH}$ for the full dataset including 7339 individuals with imputed genotypes. The orange dashed lines represent the medians. High-density genotyped individuals were chosen to be maximally unrelated and to maximise genetic diversity, which likely explains the missing right skew (i.e. a lack of inbred individuals) in their distribution (see Methods).

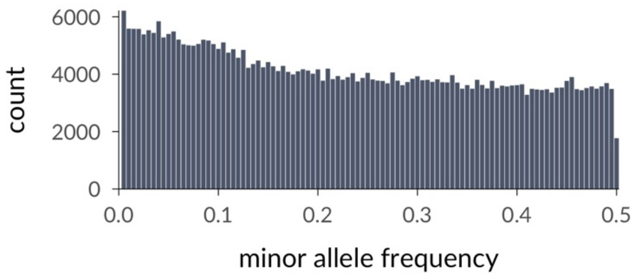


**Supplementary Figure 2: Minor allele frequency distribution.** Shown is the minor allele frequency (MAF) distribution across 417,373 imputed SNPs in 5,952 Soay sheep.

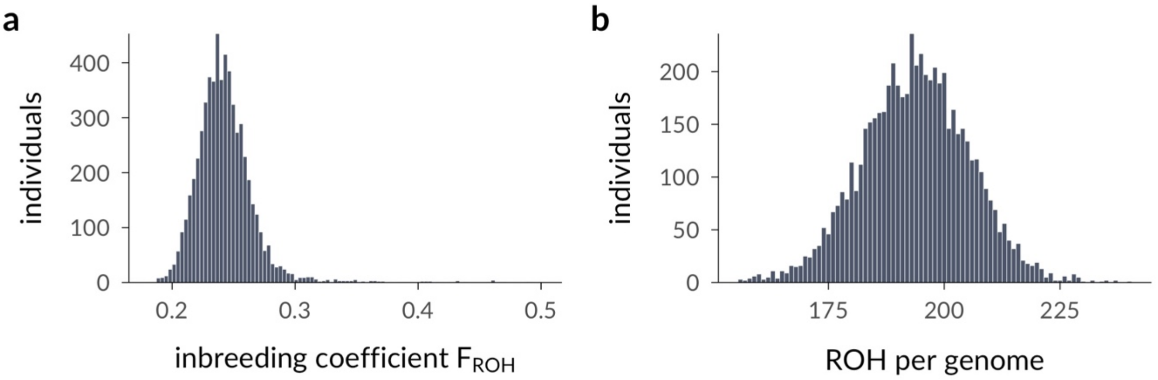


**Supplementary Figure 3: Distribution of $F_{ROH}$ and ROH. a** Distribution of inbreeding coefficients $F_{ROH}$ for 5952 individuals included in the survival analyses. **b** Distribution of the number of ROH per genome.

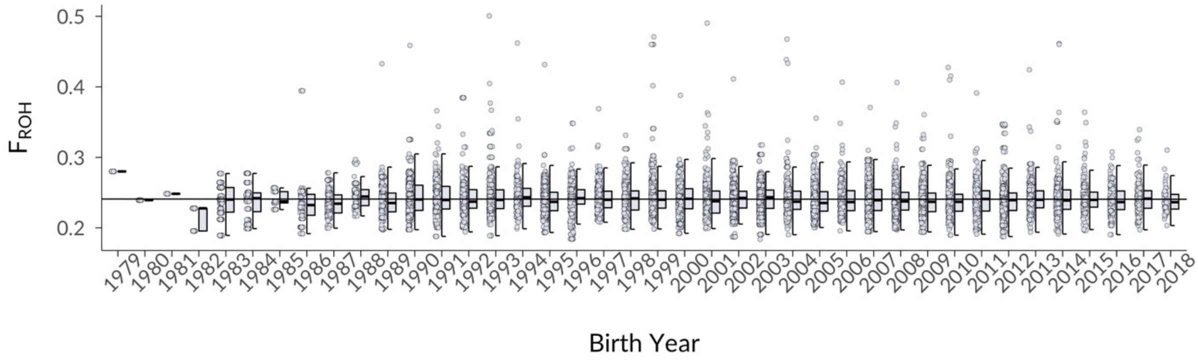


**Supplementary Figure 4: $F_{ROH}$ of individuals per birth year.** The average inbreeding coefficient did not change over the course of the study period ($\beta$ = 0, 95% CI [0,0], $p$ = 0.588 for a linear model with $F_{ROH}$ as response and birth year as numeric predictor). The plot shows the $F_{ROH}$ of individuals as points, Tukey-boxplots per birth year to sum up the distribution of $F_{ROH}$ in each year and the regression line of the linear model. Sample sizes n across 40 year (from 1979 to 2018) are 1979: 13, 1980: 15, 1981: 8, 1982: 23, 1983: 165, 1984: 164, 1985: 40, 1986: 137, 1987: 255, 1988: 145, 1989: 312, 1990: 656, 1991: 380, 1992: 532, 1993: 616, 1994: 279, 1995: 721, 1996: 464, 1997: 440, 1998: 299, 1999: 590, 2000: 626, 2001: 342, 2002: 608, 2003: 960, 2004: 359, 2005: 507, 2006: 463, 2007: 703, 2008: 703, 2009: 613, 2010: 435, 2011: 361, 2012: 501, 2013: 395, 2014: 337, 2015: 245, 2016: 230, 2017: 180, 2018: 54. The boxplots are standard Tukey boxplots (centre line = median, bounds of box = 25th and 75th percentiles, upper and lower whiskers = largest and smallest value but no further than 1.5 * inter-quartile range from the hinge).

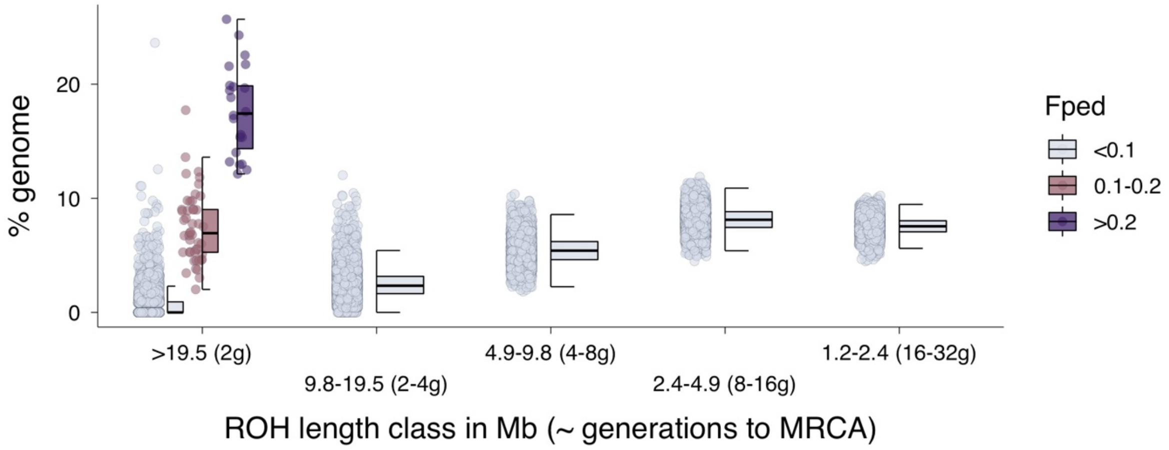

**Supplementary Figure 5: Pedigree inbreeding among individuals with ROH > 19.5Mb.** The figure shows the same data as Figure 1B with additional details on the pedigree inbreeding coefficient ($F_{ped}$) of individuals with long ROH > 19.5 Mb. $F_{ped}$ was clustered into three classes comprising more outbred individuals ($F_{ped}$ <= 0.1), moderately inbred individuals (0.1 < $F_{ped}$ < 0.2) and strongly inbred individuals ($F_{ped}$ >= 0.2). Most individuals with a large proportion of the genome in long ROH are inbred according to the pedigree.

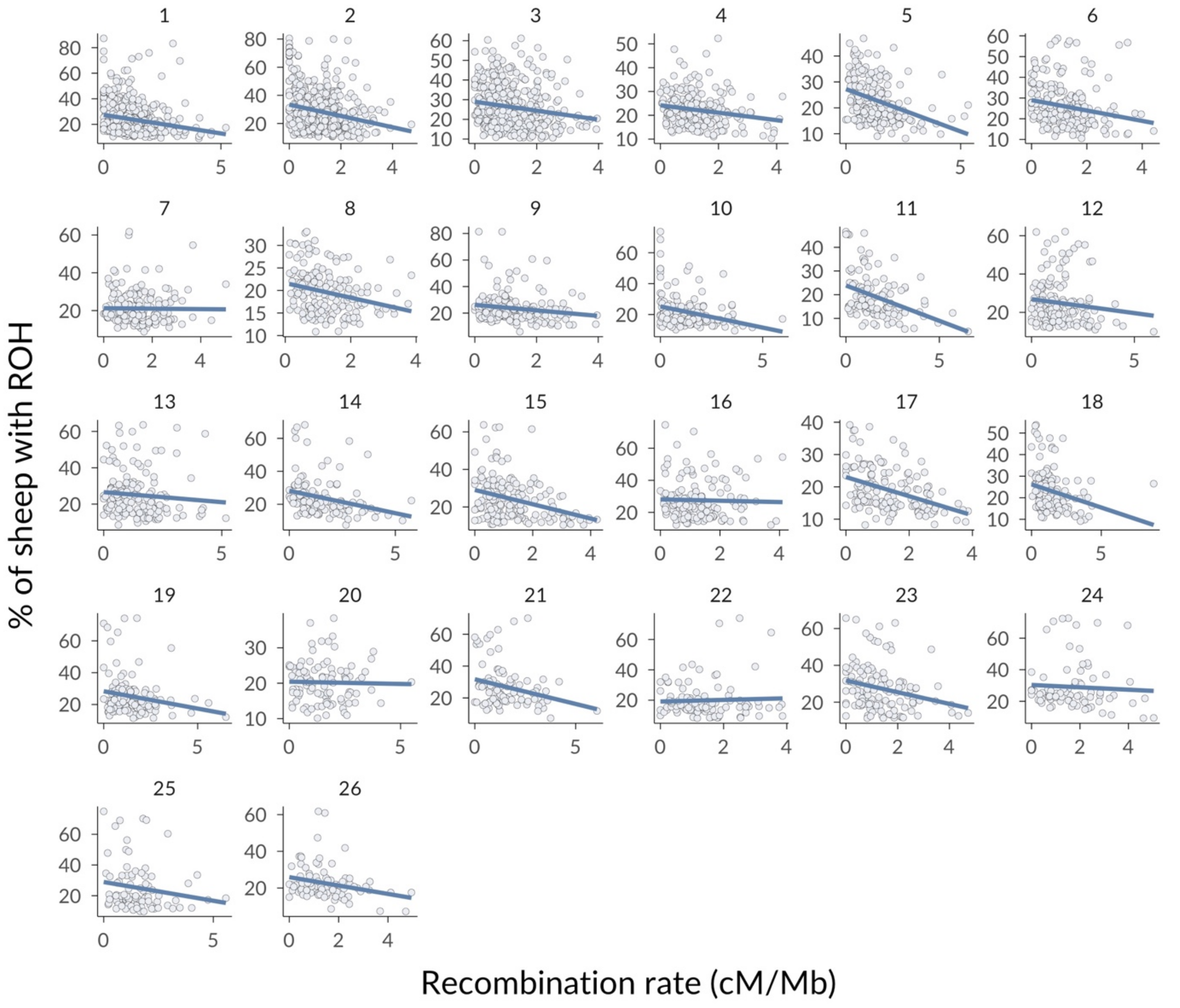


**Supplementary Figure 6: ROH frequency and recombination rate in non-overlapping 500Kb windows across the genome.** Shown is the mean frequency of ROH overlapping SNPs in 500Kb windows plotted against the rate of recombination in each window, facetted by chromosome, with linear model regression lines.

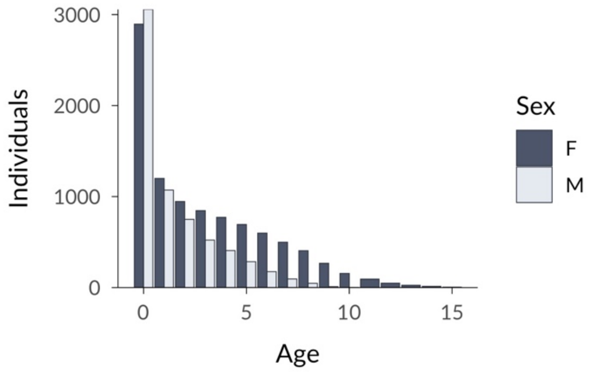


**Supplementary Figure 7: Number of annual survival observations (individuals) per age class.** Overall, the dataset consisted of 15889 observations from 5952 sheep.

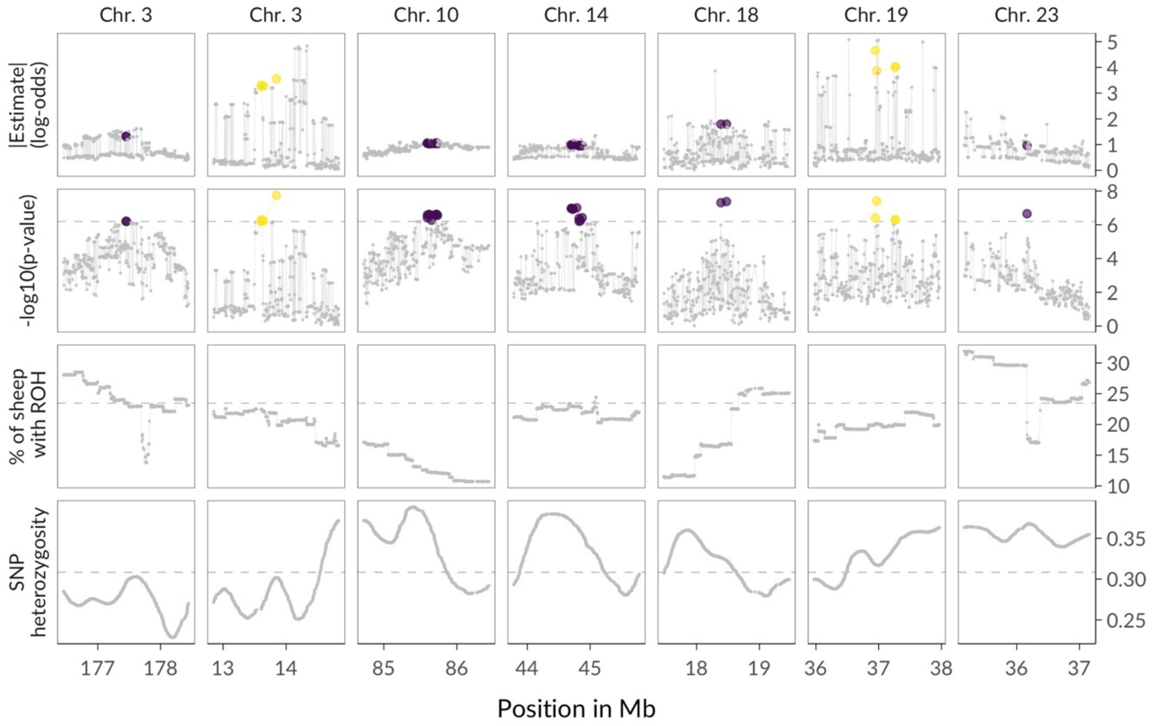


**Supplementary Figure 8: GWAS estimates and genetic diversity around top GWAS hits.** The first two rows show the GWAS model estimates (absolute) and p-values for all SNPs within a 2Mb window around the top SNP on each of the sevem genome-wide significant peaks located on six different chromosomes (see Figure 3). Purple points mark genome-wide significant negative ROH effects on survival, and yellow points mark genome-wide significant positive ROH effects on survival. Row three shows the proportion of individuals with ROH at each SNP position. Row four shows SNP heterozygosity which was smoothed using a Nadaraya-Watson kernel regression with a bandwidth of 100 SNPs. The dashed grey lines show the genome-wide significance level for p-values, and the genome-wide mean for the proportion of sheep with ROH and SNP heterozygosity.

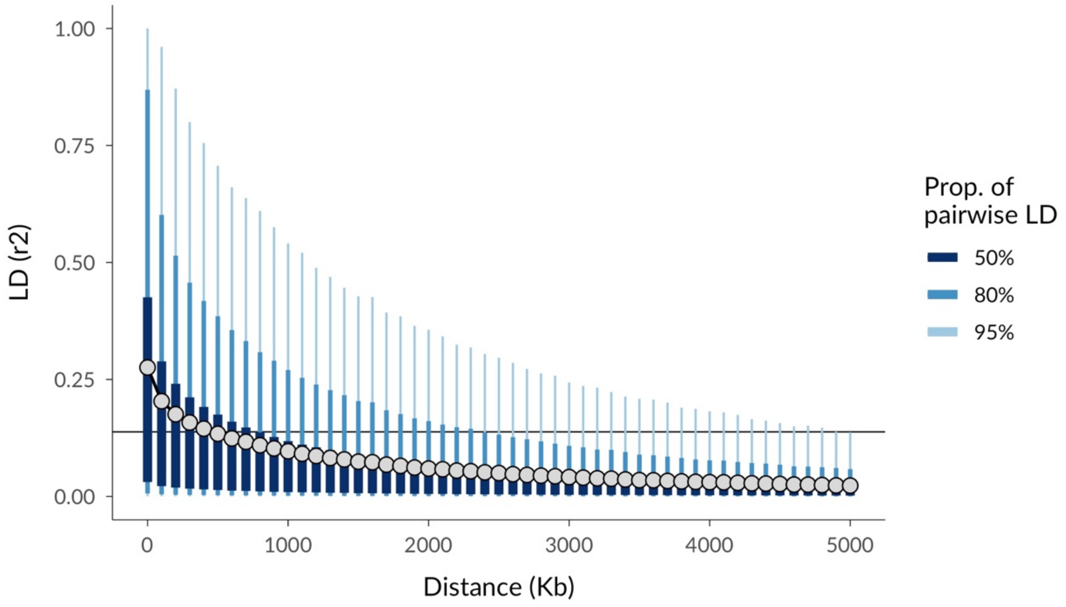


**Supplementary Figure 9: LD decay in Soay sheep.** LD was calculated based on (non-imputed) SNPs from the 50K SNP chip for all pairs of loci within each chromosome using –r2 in PLINK. The average of pairwise LD values was then calculated for loci pairs with increasing physical distance, in 100 Kb increments. The plot shows the distribution of LD $r^2$ values for loci up to 5 Mb apart. The proportion of pairwise LD comparisons per category is visualised using different shades of blue, with dark blue representing the interquartile range or middle 50% of data, and the lighter blue colors showing 80% and 95% of the data, respectively. The horizontal line is the indicator for the LD half decay. Consequently, LD half decays for loci approximately 600Kb apart (where the $6^{th}$ mean LD point falls below the line).

### Supplementary Tables

| Chromosome | % SNPs correct | % SNPs imputed | N (Snps) |
|---|---|---|---|
| 1 | 99.574 | 98.413 | 47318 |
| 2 | 99.547 | 99.180 | 41917 |
| 3 | 99.593 | 98.899 | 38265 |
| 4 | 99.454 | 99.440 | 20659 |
| 5 | 99.454 | 99.243 | 18545 |
| 6 | 99.315 | 99.584 | 19205 |
| 7 | 99.312 | 99.535 | 17647 |
| 8 | 99.726 | 99.707 | 15815 |
| 9 | 99.068 | 99.279 | 16517 |
| 10 | 99.342 | 99.541 | 15517 |
| 11 | 98.996 | 99.481 | 11567 |
| 12 | 99.420 | 99.708 | 13909 |
| 13 | 99.346 | 99.677 | 13626 |
| 14 | 98.979 | 99.425 | 10785 |
| 15 | 99.368 | 99.789 | 13897 |
| 16 | 99.528 | 99.655 | 11992 |
| 17 | 99.329 | 99.577 | 12598 |
| 18 | 98.904 | 99.467 | 11718 |
| 19 | 99.447 | 99.757 | 10230 |
| 20 | 99.109 | 99.609 | 9074 |
| 21 | 98.403 | 99.604 | 7879 |
| 22 | 99.007 | 99.706 | 9352 |
| 23 | 99.345 | 99.565 | 10004 |
| 24 | 98.628 | 99.660 | 6362 |
| 25 | 98.986 | 99.690 | 7530 |
| 26 | 99.152 | 99.614 | 7353 |

**Supplementary Table 1: Cross-validation results for genotype imputation.** To evaluate the accuracy of the genotype imputation, we masked genotypes from individuals which have been genotyped on the high-density SNP chip and used imputation to predict the masked genotypes. The table shows, for each chromosome, data summarised over the 10 cross-validation runs, in which genotypes for a single randomly chosen individual were masked and imputed per run. The second column shows the proportion of SNPs which were correctly imputed and column three shows the proportion of SNPs from the high-density array which could be imputed at all. The fourth column, N(SNPs), shows the number of SNPs on each chromosome.

| Top 0.5% ROH deserts<br>ROH density measured in 500Kb running windows | | | | |
|---|---|---|---|---|
| Chromosome | WinStart | WinEnd | % of individuals with ROH | N (SNPs) |
| 11 | 58.5 | 59.0 | 4.45 | 103 |
| 9 | 0.0 | 0.5 | 5.02 | 13 |
| 21 | 49.0 | 49.5 | 5.39 | 26 |
| 11 | 58.0 | 58.5 | 5.50 | 110 |
| 11 | 59.0 | 59.5 | 5.78 | 85 |
| 9 | 0.5 | 1.0 | 5.96 | 93 |
| 11 | 61.5 | 62.0 | 6.79 | 105 |
| 11 | 57.5 | 58.0 | 6.95 | 94 |
| 26 | 37.5 | 38.0 | 7.23 | 95 |
| 21 | 48.5 | 49.0 | 7.23 | 48 |
| 26 | 37.0 | 37.5 | 7.40 | 114 |
| 11 | 57.0 | 57.5 | 7.41 | 116 |
| 14 | 5.5 | 6.0 | 7.45 | 110 |
| 11 | 59.5 | 60.0 | 7.53 | 101 |
| 11 | 60.0 | 60.5 | 7.72 | 148 |
| 11 | 56.5 | 57.0 | 8.06 | 89 |
| 11 | 61.0 | 61.5 | 8.09 | 108 |
| 22 | 9.5 | 10.0 | 8.10 | 101 |
| 5 | 5.0 | 5.5 | 8.18 | 96 |
| 11 | 60.5 | 61.0 | 8.20 | 118 |
| 17 | 71.5 | 72.0 | 8.21 | 75 |
| 17 | 64.0 | 64.5 | 8.38 | 98 |
| 22 | 12.0 | 12.5 | 8.38 | 115 |
| 22 | 11.5 | 12.0 | 8.39 | 89 |

| Top 0.5% ROH islands<br>ROH density measured in 500Kb running windows | | | | |
|---|---|---|---|---|
| Chromosome | WinStart | WinEnd | % of individuals with ROH | N (SNPs) |
| 1 | 227.0 | 227.5 | 87.28 | 41 |
| 2 | 112.0 | 112.5 | 85.97 | 11 |
| 1 | 227.5 | 228.0 | 83.28 | 65 |
| 9 | 77.5 | 78.0 | 81.40 | 43 |
| 9 | 78.0 | 78.5 | 81.31 | 67 |
| 2 | 104.0 | 104.5 | 80.49 | 42 |
| 2 | 103.0 | 103.5 | 79.96 | 104 |
| 2 | 103.5 | 104.0 | 78.99 | 67 |
| 2 | 112.5 | 113.0 | 78.73 | 38 |
| 1 | 226.5 | 227.0 | 77.17 | 69 |
| 2 | 111.0 | 111.5 | 76.91 | 58 |
| 1 | 232.5 | 233.0 | 75.94 | 65 |
| 25 | 21.5 | 22.0 | 74.91 | 65 |
| 16 | 54.0 | 54.5 | 74.58 | 69 |
| 19 | 1.5 | 2.0 | 74.26 | 65 |
| 19 | 1.0 | 1.5 | 74.14 | 67 |
| 22 | 46.0 | 46.5 | 74.07 | 95 |
| 2 | 113.0 | 113.5 | 73.95 | 60 |
| 2 | 113.5 | 114.0 | 73.92 | 61 |
| 2 | 114.0 | 114.5 | 73.89 | 42 |
| 10 | 43.0 | 43.5 | 73.63 | 57 |
| 1 | 232.0 | 232.5 | 72.98 | 76 |
| 24 | 34.0 | 34.5 | 72.69 | 70 |
| 24 | 36.5 | 37.0 | 72.40 | 53 |

**Supplementary Table 2: Top ROH deserts and islands.** Shown are the top 0.5 % 500Kb regions with the lowest ROH density in our sample of 5952 sheep to the left and the top 0.5% 500Kb regions with the highest ROH density to the right. The start (WinStart) and end (WinEnd) of each window at the respective chromosome are given in Megabasepairs (Mb). The percentage of individuals with ROH was calculated as the average number of ROH overlapping the SNPs in a window. N (SNPs) shows how many SNPs were in a given window.

**A**

| Term | Estimate | Std.Error | CI (2.5%) | CI (97.5%) | Standardization | Info | R2 |
|---|---|---|---|---|---|---|---|
| Fixed effects | | | | | | | |
| Intercept | 24.229 | 0.481 | 23.282 | 25.191 | | | |
| Recombination rate (cM/Mb) | -2.307 | 0.121 | -2.546 | -2.063 | (x-mean(x))/sd(x) | continuous | 0.04, 95%CI [0.02, 0.07] |
| Heterozygosity | -6.97 | 0.121 | -7.199 | -6.744 | (x-mean(x))/sd(x) | continuous | 0.38, 95%CI [0.36, 0.40] |
| Random effects (variances) | | | | | | | |
| Chromosome | 2.352 | 0.37 | 1.618 | 3.097 | | n = 26 | |
| Residual | 8.307 | 0.08 | 8.144 | 8.464 | | | |

**B**

| Term | Estimate | Std.Error | CI (2.5%) | CI (97.5%) | Standardization | Info | R2 |
|---|---|---|---|---|---|---|---|
| Fixed effects | | | | | | | |
| Intercept | 31.677 | 0.532 | 30.545 | 32.892 | | | |
| Recombination rate (cM/Mb) | -1.216 | 0.12 | -1.479 | -0.962 | (x-mean(x))/sd(x) | continuous | 0.01, 95%CI [0.00, 0.04] |
| Heterozygosity | -9.189 | 0.12 | -9.439 | -8.963 | (x-mean(x))/sd(x) | continuous | 0.53, 95%CI [0.50, 0.55] |
| Random effects (variances) | | | | | | | |
| Chromosome | 2.623 | 0.38 | 1.834 | 3.353 | | n = 26 | |
| Residual | 8.269 | 0.083 | 8.11 | 8.443 | | | |

**C**

| Term | Estimate | Std.Error | CI (2.5%) | CI (97.5%) | Standardization | Info | R2 |
|---|---|---|---|---|---|---|---|
| Fixed effects | | | | | | | |
| Intercept | 13.752 | 0.355 | 13.169 | 14.372 | | | |
| Recombination rate (cM/Mb) | -2.353 | 0.101 | -2.572 | -2.165 | (x-mean(x))/sd(x) | continuous | 0.08, 95%CI [0.06, 0.11] |
| Heterozygosity | -3.395 | 0.101 | -3.585 | -3.202 | (x-mean(x))/sd(x) | continuous | 0.17, 95%CI [0.15, 0.19] |
| Random effects (variances) | | | | | | | |
| Chromosome | 1.717 | 0.256 | 1.185 | 2.208 | | n = 26 | |
| Residual | 6.912 | 0.068 | 6.774 | 7.046 | | | |

**Supplementary Table 3: Model estimates for a Gaussian mixed model of ROH prevalence in 500Kb windows across the genome.** Table A shows the results for ROH > 1.2 Mb as used for the main analyses, table B shows the results for ROH > 0.3 Mb and table C shows the results for ROH > 3 Mb. Each table presents the model estimates of a mixed model with ROH prevalence as response, recombination rate and heterozygosity as fixed effects and chromosome as random effect. ROH prevalence has been estimated as the mean ROH per non-overlapping 500Kb window divided by the number of individuals. The last column reports the marginal $R^2$ of the model (row: Fixed effects) as well as the variance explained by recombination rate and heterozygosity (reported as semi-partial $R^2$, see Methods for details).

| Term | Post.Mean | Std.Error | CI (2.5%) | CI (97.5%) | Standardisation | Info |
|---|---|---|---|---|---|---|
| Fixed effects | | | | | | |
| Intercept | 0.53 (1.7) | 0.44 (1.55) | -0.32 (0.73) | 1.42 (4.14) | | |
| $F_{ROH}$ | -0.91 (0.4) | 0.14 (1.15) | -1.2 (0.3) | -0.63 (0.53) | (x * 10)-mean(x * 10) | continuous |
| LifeStage: EarlyLife | 2.82 (16.78) | 0.08 (1.08) | 2.66 (14.3) | 3 (20.09) | | categorical (0=no, 1=yes) |
| LifeStage: MidLife | 3.61 (36.97) | 0.13 (1.14) | 3.38 (29.37) | 3.91 (49.9) | | categorical (0=no, 1=yes) |
| LifeStage: LateLife | 2.41 (11.13) | 0.19 (1.21) | 2.14 (8.5) | 2.91 (18.36) | | categorical (0=no, 1=yes) |
| Sex | -0.56 (0.57) | 0.05 (1.05) | -0.66 (0.52) | -0.46 (0.63) | | categorical (0=female, 1=male) |
| Twin | -0.74 (0.48) | 0.07 (1.07) | -0.88 (0.41) | -0.61 (0.54) | | categorical (0=no, 1=yes) |
| $F_{ROH}$ * (LifeStage: EarlyLife) | -0.21 (0.81) | 0.26 (1.3) | -0.72 (0.49) | 0.3 (1.35) | | |
| $F_{ROH}$ * (LifeStage: MidLife) | 0.12 (1.13) | 0.38 (1.46) | -0.62 (0.54) | 0.88 (2.41) | | |
| $F_{ROH}$ * (LifeStage: LateLife) | 0.5 (1.65) | 0.28 (1.32) | -0.05 (0.95) | 1.05 (2.86) | | |
| Random effects (variances) | | | | | | |
| Birth year | 0.79 | 0.18 | 0.59 | 1.24 | | n = 40 |
| Capture year | 2.57 | 0.17 | 2.36 | 2.97 | | n = 40 |
| Individual | 0.3 | 0.01 | 0.29 | 0.33 | | n = 5952 |
| Add. genetic | 0.3 | 0.02 | 0.28 | 0.34 | | Pedigree-based |

**Supplementary Table 4: Model estimates for the Bayesian binomial animal model of annual survival.** Shown are the posterior mean, standard error, lower and upper credible interval on the logit (log-odds) scale with the exponentiated estimates (odds-ratios) in round brackets, and information about the standardisation of the variables. Life stage was fitted as a factor with four levels, lamb (age = 0, reference level), early life (age = 1,2), mid life (age = 3,4), late life (5+). The last column shows whether variables were fitted as continuous or categorical and how the levels for categorical variables were coded. For the random intercept effects of birth year, capture year and individual, the last column shows the number of groups. The last row in the table shows the estimates for the additive genetic variance, based on a pedigree-derived relationship matrix. The dataset underlying the model contained 15889 observations from 5952 individuals.

| SNP | Chromosome | Position (Bp) | ROH status | Estimate (log-odds) | p-value | Allele1 | Allele2 |
|---|---|---|---|---|---|---|---|
| oar3_OAR3_177451112 | 3 | 177451112 | AA | -1.32 | $6.25 \times 10^{-7}$ | A | G |
| oar3_OAR3_13845652 | 3 | 13845652 | GG | 3.56 | $1.82 \times 10^{-8}$ | A | G |
| oar3_OAR10_85720912 | 10 | 85720912 | AA | -1.06 | $2.43 \times 10^{-7}$ | A | G |
| oar3_OAR14_44787854 | 14 | 44787854 | AA | -0.99 | $9.81 \times 10^{-8}$ | A | G |
| oar3_OAR18_18475667 | 18 | 18475667 | GG | -1.80 | $4.19 \times 10^{-8}$ | G | A |
| oar3_OAR19_36967641 | 19 | 36967641 | GG | 3.87 | $3.90 \times 10^{-8}$ | A | G |
| s23340.1 | 23 | 36164028 | GG | -0.96 | $2.21 \times 10^{-7}$ | G | A |

**Supplementary Table 5: SNPs with the strongest association between ROH status and annual survival at each GWAS peak.** At each SNP location,

the effects of two binary fixed effects were tested, one for whether each allele being part of an ROH. The ROH status column shows the allelic ROH status, alongside the model estimate and the associated two-sided p-values from Wald Z-tests.
